# CDK4/6 inhibitors enhance oxaliplatin efficacy in colorectal cancer with RB-dependent and tumor-selective activity in intestinal model

**DOI:** 10.64898/2026.04.15.718743

**Authors:** Alana S. Oliveira de Souza, Julia S. M. Conceição, Letícia S. Ferraz, João P. A. Delou, Bruna A. Miranda, Carla Verissimo, Mayra dos Santos Carneiro, Stevens K. Rehen, Martín H. Bonamino, Helena L. Borges

## Abstract

Although the retinoblastoma protein (pRB) is functionally inactivated by hyperphosphorylation in the majority of colorectal cancers (CRC) - with RB1 rarely mutated and even amplified at the genomic level - three critical gaps remain unaddressed: no study has systematically compared which first-line chemotherapeutic agent best synergizes with CDK4/6 inhibition using head-to-head quantitative analysis; functional differences between palbociclib and abemaciclib in chemotherapy combinations have not been characterized in CRC; and direct genetic evidence of RB dependency in this combinatorial context is lacking. Here, we addressed these gaps by evaluating palbociclib and abemaciclib combined with oxaliplatin, 5-fluorouracil, and SN-38 in HCT116 CRC cells, with validation in SW480 cells, RB1-silenced HCT116 cells (shRNA-RB), and non-tumoral intestinal epithelial cells (IEC-6), using quantitative drug interaction analysis (Chou-Talalay), cell cycle profiling, apoptosis assessment, and pRB phosphorylation measurement. Oxaliplatin was the most consistently synergistic partner for both CDK4/6 inhibitors (CI < 1 across all tested concentrations), while combinations with SN-38 yielded variable results and 5-FU combinations approached additivity. The oxaliplatin combination reinforced G1 arrest and enhanced cell death, with abemaciclib producing more pronounced apoptotic induction than palbociclib - an effect not explained by differential pRB target engagement (both inhibitors reduced pRB Ser807/811 phosphorylation by ∼50%), likely reflecting abemaciclib’s broader kinase inhibitory profile. shRNA-mediated RB1 silencing partially attenuated the combinatorial effect, providing direct genetic evidence that the synergy is RB-dependent. Importantly, the combination did not significantly potentiate oxaliplatin cytotoxicity in non-tumoral IEC-6 intestinal epithelial cells, in contrast to the pronounced enhancement observed in tumor cells, and synergistic benefit was preserved at sub-cytotoxic inhibitor concentrations. These findings identify oxaliplatin as the optimal chemotherapeutic partner for CDK4/6 inhibition in CRC, with a mechanism involving RB-dependent potentiation of apoptosis that is preferentially active against tumor cells and maintained at clinically relevant inhibitor doses.

## Introduction

Colorectal cancer (CRC) is the third most commonly diagnosed cancer worldwide, accounting for approximately 930,000 deaths in 2020 and representing the second leading cause of cancer-related mortality after lung cancer [1]. The overall 5-year survival rate is approximately 60–65%, but drops to 10–15% for stage IV patients, with nearly 60% developing liver metastasis [2,3]. Treatment strategies for advanced CRC rely on first-line chemotherapy regimens based on oxaliplatin, irinotecan, and 5-fluorouracil (5-FU), often combined as FOLFOX or FOLFIRI [4]. Oxaliplatin, a platinum-based agent, forms DNA cross-links that inhibit replication and induce cell cycle arrest [5]; however, its clinical use is limited by significant toxicities, including myelosuppression, gastrointestinal effects, and peripheral neuropathy [6]. Strategies that enhance oxaliplatin efficacy while enabling dose reduction could therefore improve the therapeutic index in CRC.

Uncontrolled proliferation is a hallmark of cancer, and the CDK4/6–cyclin D–pRB axis is a central regulator of the G1/S cell cycle transition [7]. CDK4/6 inhibitors such as palbociclib and abemaciclib prevent pRB phosphorylation, maintaining its tumor-suppressive function and arresting cells in G1 [8,9]. These agents are FDA-approved for hormone receptor-positive, HER2-negative breast cancer and have demonstrated preclinical activity in multiple solid tumors, including glioblastoma [10,11], pancreatic cancer [12], esophageal adenocarcinoma [13] and gastric cancer [14]. In CRC, although the CDK4/6–RB pathway is frequently disrupted across cancer types through cyclin D amplification, CDKN2A deletion, or RB1 loss [15], RB1 mutations are rare (10–15%) and the vast majority of CRC tumors retain— and even amplify — the RB1 gene [16,17]. Consistent with this, proteogenomic analysis of human colon cancer revealed that RB1 protein is overexpressed in tumors compared to adjacent normal tissue, with functional inactivation driven not by genetic loss but by hyperphosphorylation of pRB, which in turn promotes both proliferation and resistance to apoptosis [18]. This molecular context — intact RB1 with aberrant pRB phosphorylation—provides a strong biological rationale for pharmacological CDK4/6 inhibition in CRC.

Preclinical studies have begun to explore this rationale. CDK4/6 inhibitors as single agents have shown consistent activity in CRC models both in vitro and in vivo [19,20]. Schneider et al. [21] recently demonstrated that ribociclib is effective across a broad panel of CRC cell lines, including FOLFOX-resistant cells, while ribociclib-resistant cells exhibit marked reduction of RB and phospho-RB protein levels, underscoring RB status as a key determinant of CDK4/6 inhibitor response; however, no synergism was observed when ribociclib was combined with FOLFOX in that study. Regarding combination approaches with other CDK4/6 inhibitors, Zhang et al. [22] showed that palbociclib synergizes with irinotecan in CRC cells under hypoxic conditions. Yu et al. [23] identified a noncanonical role of RB1 in modulating chromatin activity through the TEAD4/HDAC1 complex, demonstrating that CDK4/6 inhibition suppresses DNA repair gene expression and sensitizes CRC to oxaliplatin through a synthetic lethality mechanism. Furthermore, the concept that CDK4/6 inhibitors can simultaneously potentiate antitumor efficacy in tumor cells while protecting normal proliferating cells has been demonstrated with trilaciclib in chemotherapy regimens for small cell lung cancer [24], supporting the notion of a CDK4/6-dependent therapeutic window.

Despite these advances, four critical gaps remain unaddressed. First, no study has performed a head-to-head comparison of oxaliplatin, 5-FU, and irinotecan as combinatorial partners for CDK4/6 inhibition in CRC using quantitative synergism analysis; published work evaluates isolated drug pairs without systematic comparison. Second, functional differences between abemaciclib and palbociclib in chemotherapy combinations have not been characterized in CRC, limiting rational inhibitor selection. Third, direct genetic evidence of RB dependency in the CDK4/6 inhibitor–chemotherapy combinatorial context is lacking in CRC; Yu et al. [23] and Schneider et al. [21] provide mechanistic and correlative evidence, but neither employed genetic RB silencing to test the combinatorial response. Fourth, whether this combination exhibits selective activity against tumor cells while preserving non-tumoral intestinal epithelial cells has not been evaluated — a clinically important question given the gastrointestinal toxicity of oxaliplatin-based regimens.

Here, we systematically evaluate palbociclib and abemaciclib combined with oxaliplatin, 5-fluorouracil, and SN-38 in HCT116 CRC cells, with validation in SW480 cells, RB1-silenced HCT116 cells (shRNA-RB), and non-tumoral intestinal epithelial cells (IEC-6). Using quantitative drug interaction analysis (Chou-Talalay), cell cycle profiling, apoptosis assessment, and pRB phosphorylation measurement, we identify oxaliplatin as the optimal chemotherapeutic partner for CDK4/6 inhibition and provide direct genetic evidence, via shRNA-mediated RB1 knockdown, that the combinatorial effect is RB-dependent. We further demonstrate preferential activity against tumor cells relative to non-tumoral intestinal epithelium, supporting a favorable therapeutic window for this combination in CRC.

## Materials and Methods

### Cell Lines and Culture Conditions

The human colorectal cancer cell lines HCT116 (ATCC: CCL-247; colorectal carcinoma) and SW480 (ATCC: CCL-228; colorectal adenocarcinoma) were obtained from the Rio de Janeiro Cell Bank (Banco de Células do Rio de Janeiro, BCRJ). Both cell lines were cultured in Dulbecco’s Modified Eagle Medium: Nutrient Mixture F-12 (DMEM/F-12; Gibco, 12400024), supplemented with 10% fetal bovine serum (FBS; Gibco, 12657029) and 2 mM GlutaMAX™ (Gibco, 35050061). The non-tumoral rat intestinal epithelial cell line IEC-6 (ATCC: CRL-1592), kindly donated by Prof. José Garcia Abreu (ICB/UFRJ), was maintained in the same basal medium (DMEM/F-12 supplemented with 10% FBS) with additional insulin supplementation at a final concentration of 0.1 U/mL throughout all experimental procedures, including treatments. All cell cultures were maintained at 37°C in a humidified atmosphere containing 5% CO₂. Mycoplasma contamination testing was performed every two months by indirect fluorescence staining with Hoechst 33258. Cell line identity was verified by short tandem repeat (STR) profiling with microsatellite markers at the Firmino Torres de Castro Macromolecular Metabolism Laboratory (UFRJ). All cell lines were used at passage numbers below 20 following STR authentication. For cryopreservation, cells were frozen in liquid nitrogen in a solution containing 90% FBS and 10% dimethyl sulfoxide (DMSO).

### Reagents and Drug Preparation

The selective CDK4/6 inhibitors abemaciclib (MedChemExpress, HY-16297A) and palbociclib (MedChemExpress, HY-A0065) were used either as single agents or in combination with one of the following chemotherapeutic agents: 5-fluorouracil (5-FU; Merck, F6627), SN-38 (7-ethyl-10-hydroxycamptothecin; Cayman Chemical, 15632), and oxaliplatin (Bergamo, 1151955). Palbociclib, abemaciclib, and SN-38 were dissolved in 100% DMSO and stored as 5 mM aliquots at −80°C. 5-FU was dissolved in DMSO and stored as 385 mM aliquots at 4°C. Oxaliplatin was dissolved in sterile water for injection and stored as 12.6 mM aliquots at −20 °C. Negative control cells (untreated) were maintained in DMEM/F12 medium supplemented with 10% fetal bovine serum (FBS) without the addition of DMSO, as the highest solvent concentration expected under the experimental conditions was 0.02%, a level previously tested and shown not to affect cytotoxicity in the colorectal cancer cell lines used.

### Cell Viability Assay on Multiparametric Imaging Platform

HCT116 cells (parental, LacZ, and shRNA-RB) were seeded at 1 × 10⁴ cells/well (for 24 h assays) or 5 × 10³ cells/well (for 48 h assays) in 96-well plates containing DMEM/F-12 supplemented with 10% FBS. SW480 cells were seeded at 1.2 × 10⁴ cells/well, and IEC-6 cells at 5 × 10³ cells/well, under the same medium conditions. After an initial adhesion period of 24 h (HCT116, IEC-6) or 48 h (SW480, to allow adequate recovery after passaging), the medium was replaced with fresh medium containing the indicated drug concentrations.

For HCT116, the following concentrations were used in combination experiments: oxaliplatin at 0.4, 0.6, and 0.8 µM; abemaciclib at 300 nM (IC₅₀); and palbociclib at 400 nM (selected as the lowest concentration within the response plateau that remains below the maximum reported plasma concentration of 355 nM [25]. For SW480, concentrations were adjusted based on individual dose–response curves: oxaliplatin at 1.5 µM (IC₅₀), abemaciclib at 191 nM (IC₂₀), and palbociclib at 95 nM (IC₂₀). IEC-6 cells were treated with the same concentrations used for HCT116 (oxaliplatin 0.6 µM, abemaciclib 300 nM, palbociclib 400 nM). Treatments were maintained for 48 h in all combination experiments unless otherwise stated.

Cell viability was assessed using the LIVE/DEAD™ Viability/Cytotoxicity Kit (Invitrogen, L3224). Live cells were labeled with calcein-AM (0.4 µM) and dead cells with ethidium homodimer-1 (EthD-1; 2 µM). Nuclei were counterstained with Hoechst 33342 (1 µM). Fluorescent dyes were diluted in Tyrode’s buffer and applied after medium removal, followed by incubation for 30 min at 37 °C. After washing and buffer replacement, images were acquired using a Cytation 5 Cell Imaging Multi-Mode Reader (BioTek/Agilent) with a 4× objective. Two non-overlapping fields were captured for each well, with three technical replicate wells for each condition for each experiment. Cell viability was quantified as the percentage of calcein-AM–positive cells relative to total Hoechst-positive nuclei, using Gen5 software (BioTek/Agilent). Results were expressed as a percentage relative to untreated controls (set to 100%).

Dose–response curves for SW480 cells were generated using an Operetta CLS High-Content Analysis System (PerkinElmer), which enabled automated single-cell segmentation of Calcein-AM–positive cells even in densely clustered cultures. Due to the tendency of SW480 cells to grow in tightly adherent clusters, reliable single-cell discrimination was not achievable on the Cytation 5 system. Therefore, the effects of combination treatments in SW480 cells were assessed by flow cytometry, using cell cycle distribution and sub-G1 quantification as readouts.

### Drug Combination Analysis

The potential synergistic, antagonistic, or additive interactions between chemotherapeutic agents and CDK4/6 inhibitors in HCT116 cells were evaluated using the Chou-Talalay method (Chou, 2006; Chou, 2010). Using CompuSyn software (version 1.0, ComboSyn Inc.), the Combination Index (CI) was calculated based on the individual and combined dose–response data from the cell viability assay. CI values < 1 indicate synergism, CI = 1 indicates an additive effect, and CI > 1 indicates antagonism. The CI was calculated according to the equation: CI = (D)₁/(Dx)₁ + (D)₂/(Dx)₂, where (D) represents the dose of each drug in the combination and (Dx) represents the dose of each drug alone required to produce the same fractional effect (Chou, 2010). Inhibitory concentrations (IC₁₀, IC₂₀, IC₃₀, and IC₅₀) were determined from 48 h dose–response curves fitted in GraphPad Prism 8.4.3 using nonlinear regression (four-parameter logistic model; log[inhibitor] vs. normalized response, variable slope) and reported with 95% confidence intervals (profile likelihood).

### RB1 Gene Silencing

Stable RB1 silencing was achieved using the Sleeping Beauty transposon system, a non-viral integration approach in which the transgene is flanked by inverted terminal repeat (ITR) sequences recognized by a transposase, allowing stable genomic integration. The shRNA sequence targeting RB1 (shRB), based on a previously validated siRNA sequence (5′-GTTGATAATGCTATGTCAA-3′) [26], was cloned into the pSUPER vector under the control of the H1 promoter and subsequently subcloned into the Neo-pT3 transposon vector between the ITRs. A LacZ-targeting shRNA (target sequence: 5′-GAACGTACGCGGAATACTTCGA-3′) was used as a negative control. For each construct, four oligonucleotides encoding sense and antisense strands with BglII/HindIII-compatible overhangs were designed, annealed, and phosphorylated prior to ligation into pSUPER.

Sense and antisense oligonucleotides were annealed in Tris/NaCl buffer (50 mM/100 mM) by heating at 90 °C for 1 min followed by incubation at 37 °C for 1 h. Phosphorylation was performed with T4 Polynucleotide Kinase, and the products were ligated into pSUPER linearized with BglII and HindIII using T4 DNA ligase. The H1 promoter–shRNA cassette was excised with SpeI and ClaI, gel-purified (GelRed® staining), and inserted into the Neo-pT3 vector digested with the same enzymes. Constructs were transformed into *E. coli* DH5α, and plasmid DNA was extracted by alkaline lysis or using the QIAfilter Plasmid Maxi Kit (Qiagen). Positive colonies were confirmed by restriction enzyme digestion and sequenced using the BigDye® Terminator v3.1 kit (Applied Biosystems) with H1rev and M13fw primers. Sequence analyses were performed using SeqMan Pro™ software (DNASTAR).

Validated constructs were electroporated into HCT116 cells using the Amaxa® Nucleofector™ IIb system (program D-032) with 10 µg of Neo-pT3-LacZ or Neo-pT3-shRB vector and 4 µg of GFP-expressing plasmid to monitor transfection efficiency. Transfected cells were selected with G418 (1000 µg/mL) for 10 days, with medium changes every 3–4 days, generating the stable lines HCT116-LacZ and HCT116-shRNA-RB. RB1 knockdown efficiency was confirmed by Western blotting.

### Western Blotting

Cells were washed with ice-cold PBS and lysed in UTB buffer (75 mM Tris-HCl, 9 M urea, pH 7.4) supplemented with Pierce Protease and Phosphatase Inhibitor Mini Tablets, EDTA-free (Thermo Scientific, A32961), prepared according to the manufacturer’s instructions. Protein concentration was determined by Bradford assay using Protein Assay Dye Reagent Concentrate (Bio-Rad). Samples containing 40 µg of total protein were prepared in Laemmli sample buffer (10% SDS, 10 mM β-mercaptoethanol, 20% glycerol, 0.2 M Tris-HCl pH 6.8, bromophenol blue), denatured at 95 °C for 5 min, and resolved on 8% SDS-PAGE gels alongside a molecular weight standard (Precision Plus Protein™ Kaleidoscope™, Bio-Rad). Proteins were transferred onto polyvinylidene difluoride (PVDF) membranes, pre-activated with absolute ethanol, using a semi-dry transfer system (Bio-Rad) at 0.22 A for 1 h 20 min in transfer buffer (25 mM Tris, 192 mM glycine, 0.2% SDS, 20% ethanol).

Membranes were blocked with 3% BSA in TBS-T (150 mM NaCl, 10 mM Tris, 0.1% Tween 20) and incubated overnight (16 h) at 4 °C with the following primary antibodies: phospho-RB Ser807/811 (1:1000, Cell Signaling, 9308), total RB (1:1000, kindly donated by Dr. Jean Wang, UCSD), and α-tubulin (1:2000, Sigma, T6199). After three washes with TBS-T, membranes were incubated for 2 h at room temperature with HRP-conjugated secondary antibodies: anti-rabbit IgG (1:2000, Life Technologies, A16104) and anti-mouse IgG (1:1000, Invitrogen, G21040). Protein bands were detected by enhanced chemiluminescence (Luminata™ or Immobilon Forte™, Millipore) and visualized using the ChemiDoc™ XRS+ imaging system (Bio-Rad). Densitometric analysis was performed using Image Lab™ software (Bio-Rad), with band intensities normalized to the α-tubulin loading control and expressed relative to the untreated control condition.

### Cell Cycle Analysis by Flow Cytometry

HCT116 cells (6 × 10⁴ cells/well) and SW480 cells (7.2 × 10⁴ cells/well) were seeded in 24-well plates containing DMEM/F-12 supplemented with 10% FBS and 2 mM GlutaMAX™ and allowed to adhere for 24 h (HCT116) or 48 h (SW480, to allow adequate recovery after passaging). Cells were then treated for 48 h with oxaliplatin alone or in combination with palbociclib or abemaciclib at the concentrations specified for each cell line. After treatment, cells were trypsinized, collected in complete medium, centrifuged (2000 rpm, 5 min, 4 °C), and fixed in ice-cold 70% ethanol for at least 30 min at 4 °C. Fixed cells were washed twice with PBS, incubated with RNase A (100 µg/mL) for 15 min at room temperature, and stained with propidium iodide (PI; 10 µg/mL final concentration) diluted in PBS, protected from light.

Samples were acquired on a FACSCanto™ II flow cytometer (BD Biosciences) equipped with a 488 nm solid-state laser (Coherent® Sapphire™ 488-20), a 556 LP dichroic mirror, and a 575/26 nm bandpass filter for PI detection. A minimum of 30,000 events were collected for each sample. Data acquisition was performed using FACSDiva™ software (BD Biosciences), and cell cycle distribution was analyzed using FlowJo™ software (Tree Star Inc.). The sub-G1 population, indicative of DNA fragmentation and cell death, was quantified from the ungated PI histogram.

### Statistical Analysis

Statistical analyses were performed using GraphPad Prism version 8.4.3 (GraphPad Software, San Diego, CA, USA). Data are presented as mean ± SEM from at least three independent biological experiments, unless otherwise indicated. Each biological replicate represents an independent experiment performed on a different day. Technical replicates within each experiment were averaged before statistical analysis.

For experiments involving a single experimental factor, differences among groups were analyzed using ordinary one-way analysis of variance (ANOVA). Assumptions of normality and homogeneity of variance were assessed using GraphPad Prism. Post-hoc tests were selected according to the experimental design and biological hypotheses addressed in each experiment. Dunnett’s multiple comparison test was used when several treatments were compared against a single reference group. In experiments designed to determine whether combination treatments differed from their corresponding single-agent treatments, Dunnett’s test was applied using the combination group as the reference condition to allow direct comparison between the combination treatment and each monotherapy arm. Šídák’s test was used to control the family-wise error rate while restricting inference to predefined contrasts within treatment arms (for example, comparisons between treatments with and without CDK4/6 inhibitors within the same chemotherapeutic condition). A p value < 0.05 was considered statistically significant.

Dose-response experiments were analyzed using nonlinear regression curve fitting to estimate half-maximal inhibitory concentrations (IC₅₀), as implemented in GraphPad Prism. IC values were derived from fitted curves and reported with 95% confidence intervals (profile likelihood).

Drug combination effects were evaluated using the Combination Index (CI) method according to the Chou–Talalay model [27,28], with CompuSyn software (ComboSyn Inc., Paramus, NJ). Dose-response curves were derived from calcein-AM-based cell viability assays. CI < 1 indicates synergism, CI = 1 indicates additive effects, and CI > 1 indicates antagonism.

## Results

### Dose–response profiling identifies treatment conditions for combinatorial analysis in HCT116 cells

To establish optimal treatment conditions, we performed time-course dose–response experiments in HCT116 cells, assessing viability by Calcein AM fluorescence relative to untreated controls. Cells were treated for 24 or 48 hours with increasing concentrations of oxaliplatin, SN-38, 5-FU, palbociclib, or abemaciclib (data not shown). At 48 h, low concentrations of all three chemotherapeutic agents produced measurable reductions in viability, whereas CDK4/6 inhibitors showed comparable effects at both time points; therefore, 48 h was selected for all subsequent experiments.

Extended dose–response curves at 48 h with additional concentration points were then generated to reliably determine inhibitory concentrations (Fig. 1). For 5-FU, the IC₁₀, IC₃₀, and IC₅₀ values in HCT116 cells were 2.8, 4.5, and 6 µM, respectively (Fig. 1A). For SN-38, the corresponding values were 0.8, 1.2, and 1.6 nM (Fig. 1B), and for oxaliplatin, 0.4, 0.6, and 0.8 µM (Fig. 1C). For the CDK4/6 inhibitors, the IC₅₀ of abemaciclib was 300 nM, while palbociclib yielded an IC₅₀ of 1023 nM (Fig. 1D). Since the palbociclib dose–response curve reached a plateau encompassing three concentrations (400, 800, and 1023 nM), and 1023 nM exceeds the maximum reported plasma concentration (Cmax ∼355 nM; [25], we selected 400 nM—the lowest concentration within the plateau—as the working concentration for subsequent combination experiments.

**Figure 1.**
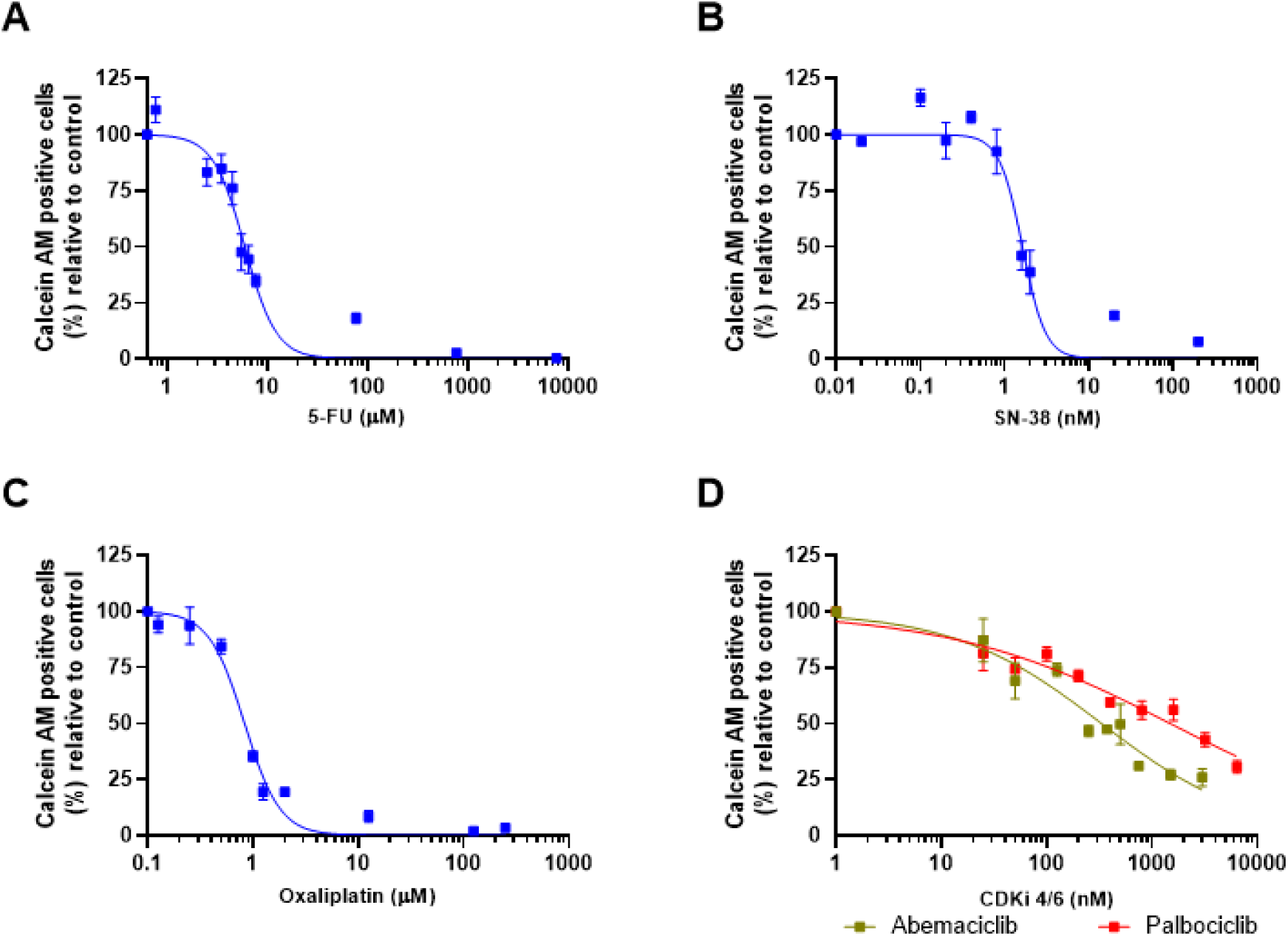
Dose–response curves of chemotherapeutic agents and CDK4/6 inhibitors in HCT116 colorectal cancer cells after 48 h of treatment. HCT116 cells (5 × 10³ cells/well) were exposed for 48 h to increasing concentrations of (A) 5-fluorouracil (0.77–7700 μM), (B) SN-38 (0.02–200 nM), (C) oxaliplatin (0.125–250 μM), and (D) the CDK4/6 inhibitors abemaciclib (25–3000 nM) or palbociclib (25–6400 nM). Cell viability was assessed by Calcein AM fluorescence and expressed as a percentage of the untreated control (100%). The data are presented as the mean ± SEM of six independent biological replicates. Dose–response curves were fitted by nonlinear regression using a four-parameter logistic model (variable slope; log[inhibitor] vs. normalized response) in GraphPad Prism 8.4.3.

### Oxaliplatin is the most synergistic chemotherapeutic partner for CDK4/6 inhibitors in HCT116 cells

To identify which chemotherapeutic agent best cooperates with CDK4/6 inhibition, HCT116 cells were treated with increasing concentrations of oxaliplatin, SN-38, or 5-FU alone or in combination with a fixed concentration of abemaciclib (300 nM) or palbociclib (400 nM), and cell viability was assessed at 48 h (Fig. 2; representative images in Fig. 3).

**Figure 2.**
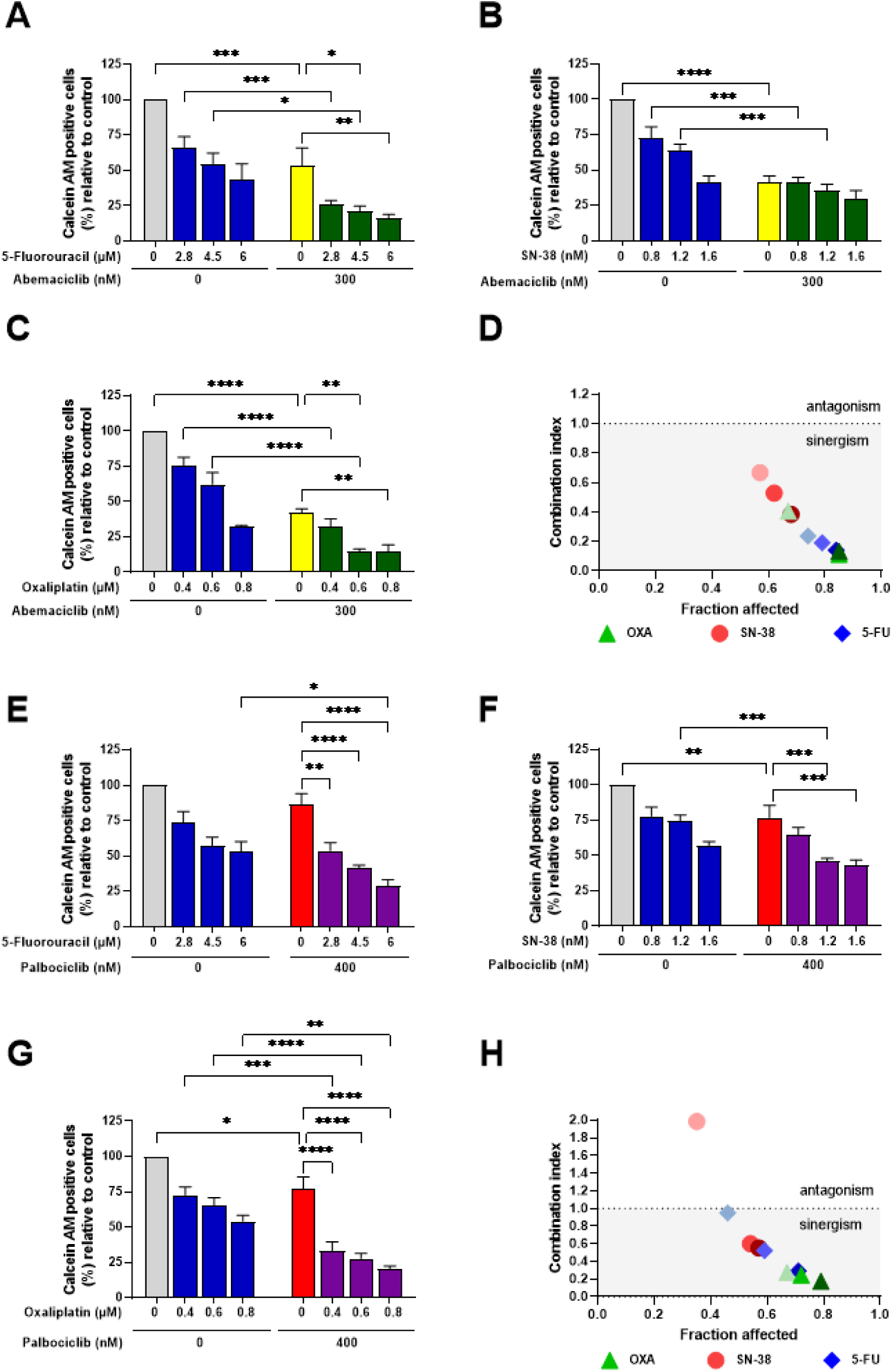
Effects of combined treatment with chemotherapeutic agents and CDK4/6 inhibitors on HCT116 cell viability and analysis of drug interaction. HCT116 cells were treated with increasing concentrations of chemotherapeutic agents in the absence or presence of fixed concentrations of CDK4/6 inhibitors, and cell viability was assessed by Calcein AM fluorescence and expressed as a percentage relative to the untreated control (100%). (A) 5-fluorouracil, (B) SN-38, (C) oxaliplatin, all of them alone or in combination with abemaciclib (300 nM). (D) Combination index (CI) analysis plotted as fraction affected versus CI for each drug combination with abemaciclib. The dashed line indicates CI = 1, where values <1 indicate synergism and values >1 indicate antagonism. (E) 5-fluorouracil, (F) SN-38, or (G) oxaliplatin, all of them alone or in combination with palbociclib (400 nM). (H) Combination index analysis plotted as fraction affected versus CI for each drug combination with palbociclib. Data are presented as mean ± SEM of at least three independent biological replicates. Bars are color-coded as follows: control (gray), chemotherapy alone (blue), abemaciclib alone (yellow), palbociclib alone (red), chemotherapy + abemaciclib (green), and chemotherapy + palbociclib (purple). Statistical significance was evaluated using ordinary one-way ANOVA followed by Šídák’s multiple comparisons test. Significance levels are indicated as follows: *p < 0.05, **p < 0.01, ***p < 0.001, ***p < 0.0001. To improve visual clarity, statistical comparisons between chemotherapy treatments alone (blue bars) and the untreated control (gray bar) are not shown in the graphs but are summarized here. ABE arm: A: CTL vs ABE ***p=0.0001; 5-FU 2.8 vs 2.8+ABE ***p=0.0005; 4.5 vs 4.5+ABE *p=0.0128; 6 vs 6+ABE ns. B: CTL vs ABE ****p<0.0001; SN-38 0.8 vs 0.8+ABE ***p=0.0009; 1.2 vs 1.2+ABE ***p=0.0004; 1.6 vs 1.6+ABE ns. C: CTL vs ABE ****p<0.0001; OXA 0.4 vs 0.4+ABE ****p<0.0001; 0.6 vs 0.6+ABE ****p<0.0001; 0.8 vs 0.8+ABE ns. PALB arm: E: CTL vs PALB ns; 5-FU 2.8 vs 2.8+PALB ns; PALB vs 2.8+PALB **p=0.0040; PALB vs 4.5+PALB ****p<0.0001; PALB vs 6+PALB ****p<0.0001; 6 vs 6+PALB *p=0.0463. F: CTL vs PALB **p=0.0071; SN-38 1.2 vs 1.2+PALB ***p=0.0008; 0.8 vs 0.8+PALB ns; 1.6 vs 1.6+PALB ns; PALB vs 1.2+PALB ***p=0.0002; PALB vs 1.6+PALB ***p=0.0003. G: CTL vs PALB *p=0.0478; OXA 0.4 vs 0.4+PALB ***p=0.0007; 0.6 vs 0.6+PALB ****p<0.0001; 0.8 vs 0.8+PALB **p=0.0041; PALB vs all OXA+PALB ****p<0.0001.

**Figure 3.**
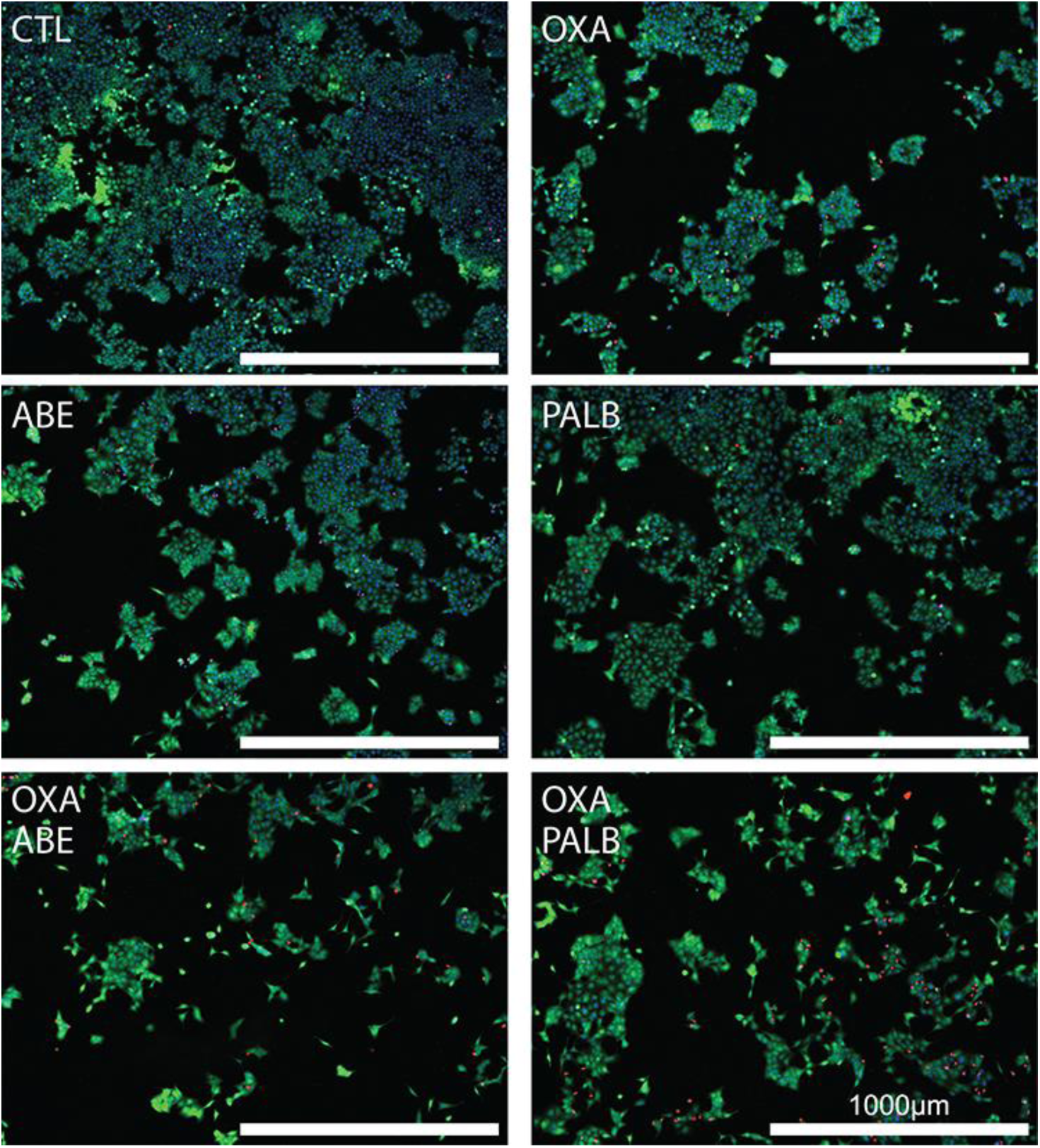
Representative fluorescence images of HCT116 cells following treatment with oxaliplatin and CDK4/6 inhibitors. HCT116 cells were treated for 48 h with oxaliplatin (0.6 μM), abemaciclib (300 nM), palbociclib (400 nM), or their respective combinations. Total cells were stained with Hoechst (blue, nuclei); viable cells with Calcein AM (green); and dead cells with Ethidium Homodimer-1 (red). Representative fields are shown for each condition: control (CTL), oxaliplatin (OXA), abemaciclib (ABE), palbociclib (PALB), oxaliplatin plus abemaciclib (OXA + ABE), and oxaliplatin plus palbociclib (OXA + PALB). Scale bar = 1000 μm.

When combined with abemaciclib, oxaliplatin at 0.6 µM significantly reduced the percentage of viable cells compared to either drug alone (Fig. 2C). Similarly, combinations of SN-38 at 0.8 and 1.2 nM with abemaciclib significantly reduced viability relative to chemotherapy alone (Fig. 2B), while 4.5 µM 5-FU combined with abemaciclib significantly reduced the percentage of viable cells compared to each drug alone (Fig. 2A). When combined with palbociclib, oxaliplatin at 0.4, 0.6, and 0.8 µM consistently and significantly reduced viability compared to monotherapies (Fig. 2G). SN-38 at 1.2 nM combined with palbociclib showed a significant effect compared to both monotherapies (Fig. 2F), whereas the effects of 5-FU–palbociclib combinations were largely limited to comparisons against the inhibitor alone (Fig. 2E).

Notably, all concentrations used were within the range of clinically achievable plasma levels. The equivalent molar concentrations derived from reported Cmax values were 3.62 µM for oxaliplatin [29], 588 nM for abemaciclib [8], and 355 nM for palbociclib [25], confirming that the experimental conditions approximate pharmacologically relevant exposures.

To quantify drug interactions, the Combination Index (CI) was calculated using the Chou-Talalay method (Fig. 2D, 2H). The analysis revealed that oxaliplatin combined with both abemaciclib and palbociclib consistently yielded CI values below 1, indicating strong synergism across all tested concentrations. Combinations with SN-38 showed variable results: while some were synergistic, others were antagonistic (e.g., SN-38 at 0.8 nM with palbociclib). Combinations with 5-FU were generally synergistic but with weaker effect sizes, approaching additivity in some conditions (e.g., 5-FU at 2.8 µM with palbociclib). Based on these results, oxaliplatin at 0.6 µM was chosen for all future mechanistic studies because it was the most consistently synergistic partner.

### Combined treatment reinforces G1 arrest and increases cell death, particularly with abemaciclib

Having identified a synergistic interaction between oxaliplatin and CDK4/6 inhibitors, we next investigated the cellular mechanisms underlying this effect. HCT116 cells were treated for 48 h with oxaliplatin (0.6 µM), abemaciclib (300 nM), palbociclib (400 nM), or their respective combinations, and cell death and cell cycle distribution were analyzed.

Quantification of ethidium homodimer-positive cells revealed a significant increase in cell death following combination treatment with oxaliplatin and abemaciclib (Fig. 4A) or palbociclib (Fig. 4B) compared to monotherapies. Flow cytometric analysis of DNA content showed that the combination of oxaliplatin with abemaciclib significantly increased the sub-G1 population, indicative of DNA fragmentation and cell death (Fig. 4C). In contrast, the combination with palbociclib did not significantly increase sub-G1 cells (Fig. 4D), suggesting that the two inhibitors differ in their capacity to induce cell death when combined with oxaliplatin.

**Figure 4.**
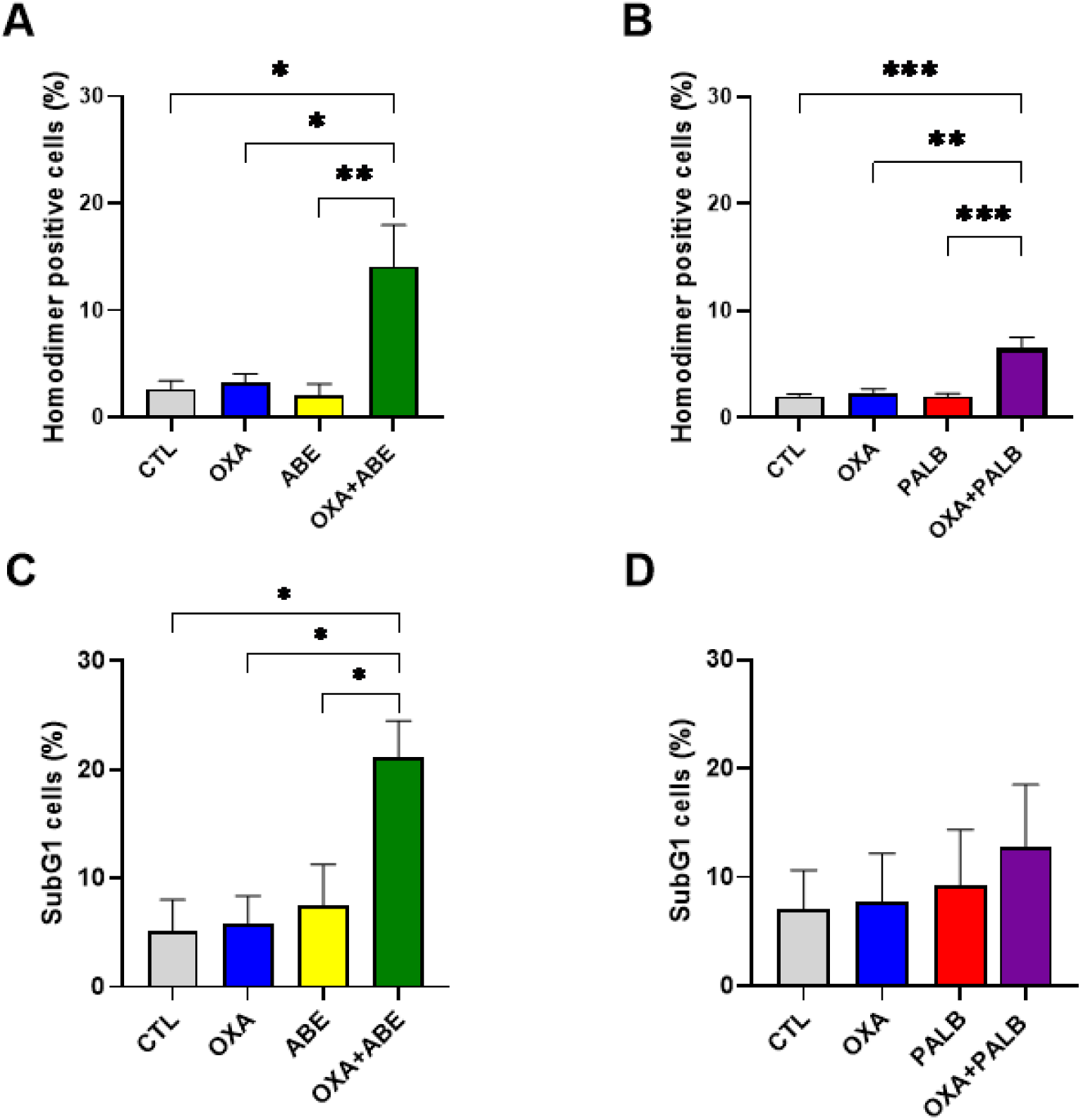
Effects of oxaliplatin and CDK4/6 inhibitors on cell death in HCT116 cells. HCT116 cells were treated for 48 h with oxaliplatin (0.6 μM), abemaciclib (300 nM), palbociclib (400 nM), or their combinations. (A-B) Cell death quantified by homodimer staining. (C–D) DNA fragmentation quantified as the SubG1 population by flow cytometry following treatment with oxaliplatin combined with abemaciclib (C) or palbociclib (D). Data are presented as the mean ± SEM of three independent biological replicates. Statistical significance was evaluated using ordinary one-way ANOVA followed by Dunnett’s multiple comparisons test. Significance levels are indicated as follows: *p < 0.05, **p < 0.01, ***p < 0.001.

Analysis of cell cycle distribution (Fig. 5A) revealed that abemaciclib as a single agent induced a robust G1 arrest, which was maintained in the presence of oxaliplatin (Fig. 5B). For palbociclib, G1 arrest was observed in the combination with oxaliplatin (Fig. 5C); although palbociclib monotherapy did not produce a marked shift in the cell cycle distribution by descriptive analysis; statistical testing confirmed a significant increase in the G1 fraction under both monotherapy and combination conditions.

**Figure 5.**
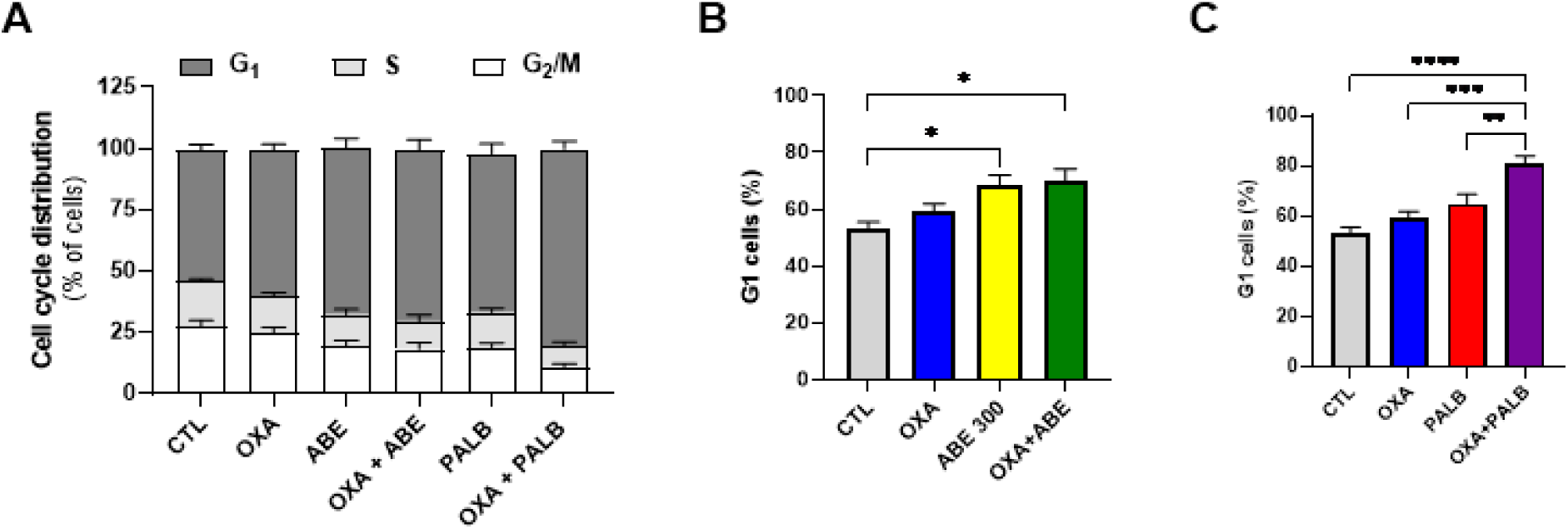
Cell cycle distribution and G1 phase quantification in HCT116 cells following treatment with oxaliplatin and CDK4/6 inhibitors. (A) Distribution of cell cycle phases (G1, S, and G2/M) in HCT116 cells treated with oxaliplatin (OXA), abemaciclib (ABE), palbociclib (PALB), or their combinations. (B) Quantification of G1 phase cells following treatment with OXA, ABE, or OXA+ABE. (C) Quantification of G1 phase cells following treatment with OXA, PALB, or OXA+PALB. The data are presented as the mean ± SEM of at least three independent biological replicates. Statistical analysis was performed using ordinary one-way ANOVA followed by Šídák’s multiple comparisons test within each treatment arm. Significance levels are indicated by asterisks as shown in the graphs. Significance levels are indicated as follows: *p < 0.05, **p < 0.01, ***p < 0.001, ****p < 0.0001.

Since CDK4/6 inhibitors exert their cell cycle effects through inhibition of pRB phosphorylation, we assessed phospho-RB (Ser807/811) levels by Western blotting. Both abemaciclib and palbociclib reduced pRB phosphorylation by approximately 50% compared to untreated controls (Fig. 6), confirming on-target activity consistent with CDK4/6 inhibition.

**Figure 6.**
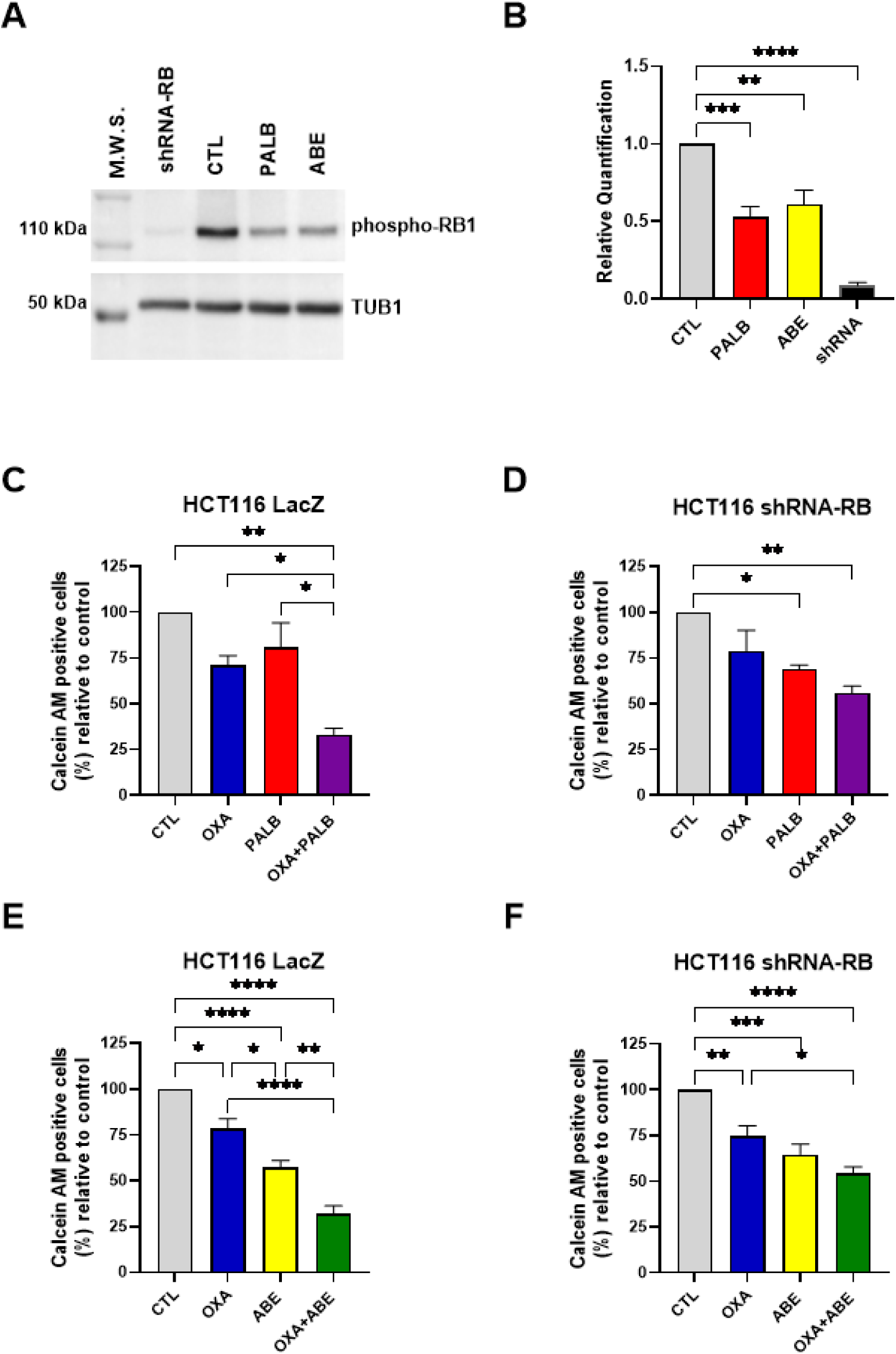
Effects of oxaliplatin combined with CDK4/6 inhibitors on cell viability in HCT116 cells with different RB1 backgrounds. (A) Representative western blot showing pRB1 expression in HCT116 whole-cell lysates (M.W.S.), RB1 knockdown (shRNA-RB), untreated control (CTL), palbociclib-treated (PALB), and abemaciclib-treated (ABE) cells. TUB1 was used as a loading control. Molecular weight markers are indicated. (B) Densitometric quantification of pRB1 normalized to TUB1 and expressed relative to control. Data are presented as the mean ± SEM of four independent biological replicates. (C-F) HCT116 LacZ control (LacZ), and RB1-silenced (shRNA-RB) cells were treated for 48 h with oxaliplatin (0.6 μM), palbociclib (400 nM), abemaciclib (300 nM), or their respective combinations. Cell viability was assessed by Calcein AM fluorescence and expressed as a percentage relative to the untreated control within each cell line (set to 100%). (C–D) Effects of oxaliplatin combined with palbociclib. (E–F) Effects of oxaliplatin combined with abemaciclib. Data are presented as mean ± SEM of three (PALB) or four (ABE) independent biological replicates. Statistical analysis was performed using ordinary one-way ANOVA followed by Šídák’s multiple comparisons test. Significance levels are indicated as follows: *p < 0.05; **p < 0.01; ***p < 0.001; ****p < 0.0001.

In summary, these findings suggest that the synergistic decrease in viability seen in HCT116 cells is due to a combination of prolonged G1 arrest and increased cell death. Abemaciclib has a stronger effect on both sub-G1 accumulation and G1 arrest than palbociclib.

### The cytotoxic effect of combined treatment is partially dependent on RB expression

To determine whether the enhanced cytotoxicity of combined treatment requires RB, we generated stable HCT116 cell lines with shRNA-mediated RB1 silencing (shRNA-RB) or a LacZ-targeting control (LacZ), based on a previously validated siRNA sequence [39]. Efficient RB1 knockdown was confirmed by total RB protein Western blotting (data not shown) and by phopho-RB (Fig. 6 A, B).

LacZ control cells and shRNA-RB cells were treated for 48 h with oxaliplatin (0.6 µM) alone or in combination with abemaciclib (300 nM) or palbociclib (400 nM) (Fig. 6). In LacZ control cells, combined treatment with either inhibitor significantly reduced cell viability compared to monotherapies (Fig. 6C, E), recapitulating the results observed in parental HCT116 cells. In contrast, in RB1-silenced cells, the viability reduction induced by the combination was partially attenuated for both palbociclib (Fig. 6D) and abemaciclib (Fig. 6F). These findings suggest that the activity of CDK4/6 inhibitors in combination with oxaliplatin depends, at least in part, on RB expression.

### Sub-cytotoxic concentrations of CDK4/6 inhibitors enhance oxaliplatin-induced cell death in SW480 cells

Dose–response curves for SW480 cells were generated using Calcein-AM fluorescence on the Operetta CLS platform, which enabled single-cell segmentation in clustered cultures (Fig. 7). However, this instrument was not available for subsequent combination experiments, and the Cytation 5 platform did not reliably discriminate individual SW480 cells due to their tightly adherent growth pattern. Therefore, the effects of combination treatments in SW480 cells were assessed by flow cytometry.

**Figure 7.**
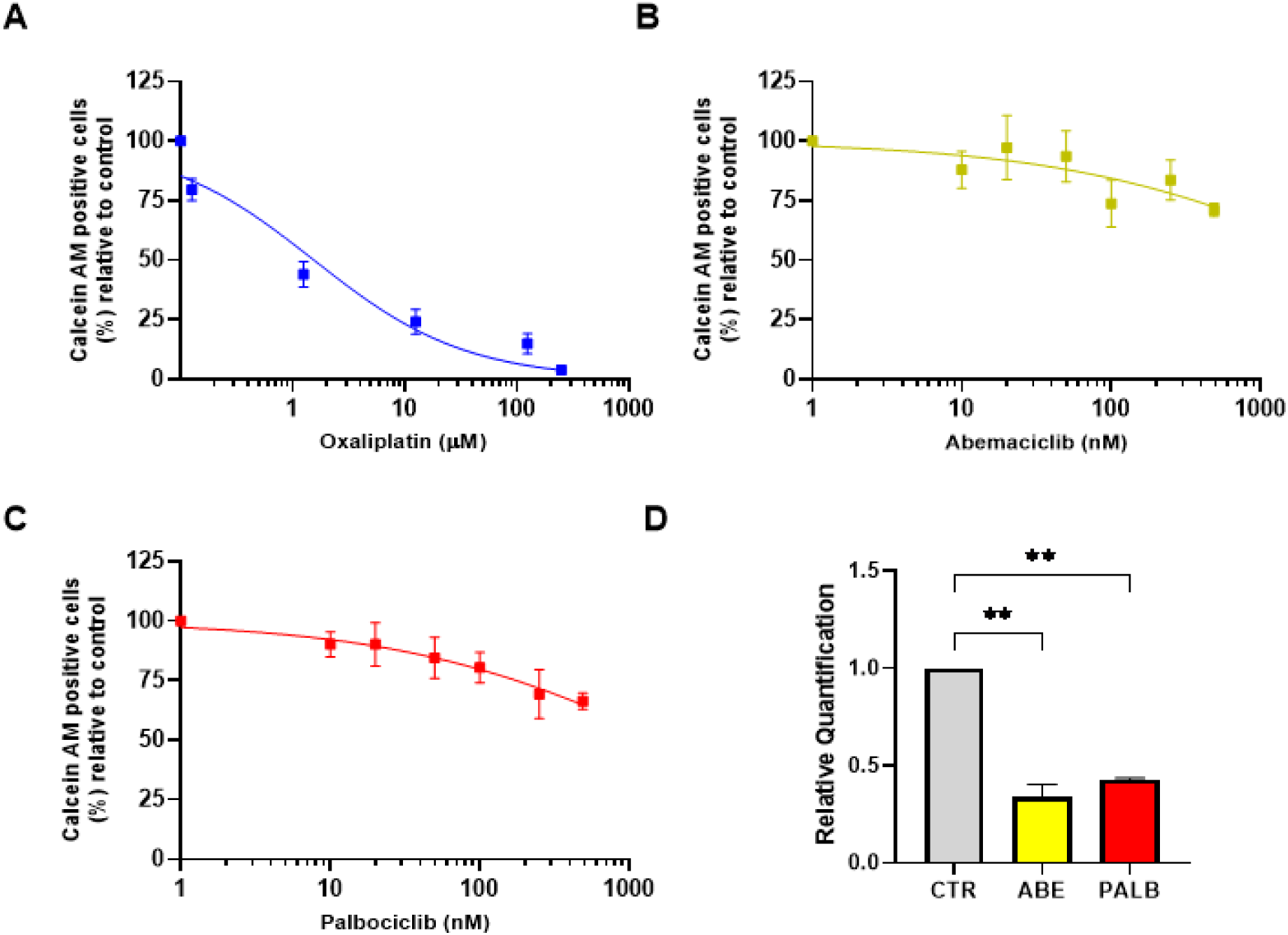
Dose–response curves of oxaliplatin and CDK4/6 inhibitors and analysis of phospho-RB levels in SW480 cells after 48 h of treatment. SW480 cells were treated for 48 h with increasing concentrations of (A) oxaliplatin (0.125–250 μM), (B) abemaciclib (10–500 nM), (C) palbociclib (10–500 nM). Cell viability was assessed by Calcein AM fluorescence and This is expressed as a percentage relative to the untreated control, which is set to 100%. The data are presented as the mean ± SEM of three independent biological replicates. Dose–response curves were fitted in GraphPad Prism using nonlinear regression (four-parameter logistic model; log[inhibitor] vs. normalized response, variable slope), and IC20 values were obtained from the fitted curves with 95% confidence intervals. (D) Densitometric quantification of pRB1 (Ser807/811) normalized to TUB1 and expressed relative to control: untreated control (CTL), abemaciclib-treated (ABE, 191 nM) cells, and palbociclib-treated (PALB, 95 nM). Data are presented as the mean ± SEM of two independent biological replicates. Statistical significance was evaluated using ordinary one-way ANOVA followed by Dunnett’s multiple comparisons test versus control. Significance levels are indicated as follows: **p < 0.01.

SW480 cells exhibited lower overall sensitivity to CDK4/6 inhibitors compared to HCT116 cells, with shallow dose–response curves that did not reach 50% inhibition within the tested concentration range (Fig. 7B, 7C). Accordingly, we determined the IC₅₀ for oxaliplatin to be 1.5 µM and the IC₂₀ values for abemaciclib and palbociclib to be 191 nM and 95 nM, respectively. Rather than pursuing higher concentrations beyond the tested range, we selected IC₂₀ doses as functionally meaningful thresholds, given that pRB target engagement was preserved at these concentrations. To confirm that these sub-cytotoxic concentrations retain on-target activity, we assessed pRB phosphorylation at Ser807/811 in SW480 cells treated with IC₂₀ doses of each inhibitor. Both palbociclib (95 nM) and abemaciclib (191 nM) reduced pRB phosphorylation by approximately 50% compared to untreated controls (Fig. 7D), indicating that CDK4/6 target engagement is preserved even at sub-cytotoxic concentrations in this cell line. This strategy tests whether sub-cytotoxic inhibitor doses—which would be more favorable in a clinical setting given the cost and toxicity profile of these agents—can potentiate the effect of conventional chemotherapy across different CRC contexts. All concentrations used remained within clinically achievable plasma levels [8, 25, 29].

SW480 cells were treated for 48 h with the indicated concentrations as monotherapies or in combination, and cell cycle distribution was analyzed by flow cytometry (Fig. 8). Abemaciclib induced a significant G1 arrest compared to control and oxaliplatin alone (Fig. 8A), whereas palbociclib did not significantly alter G1-phase distribution (Fig. 8B). The combination of oxaliplatin with abemaciclib significantly increased the sub-G1 population compared to either monotherapy (Fig. 8C), indicating enhanced cell death. The combination with palbociclib also increased the sub-G1 fraction relative to control and oxaliplatin alone (Fig. 8D). These results confirm that even at sub-cytotoxic concentrations, CDK4/6 inhibitors potentiate oxaliplatin-induced cell death across CRC cell lines, with abemaciclib additionally promoting G1 arrest.

**Figure 8.**
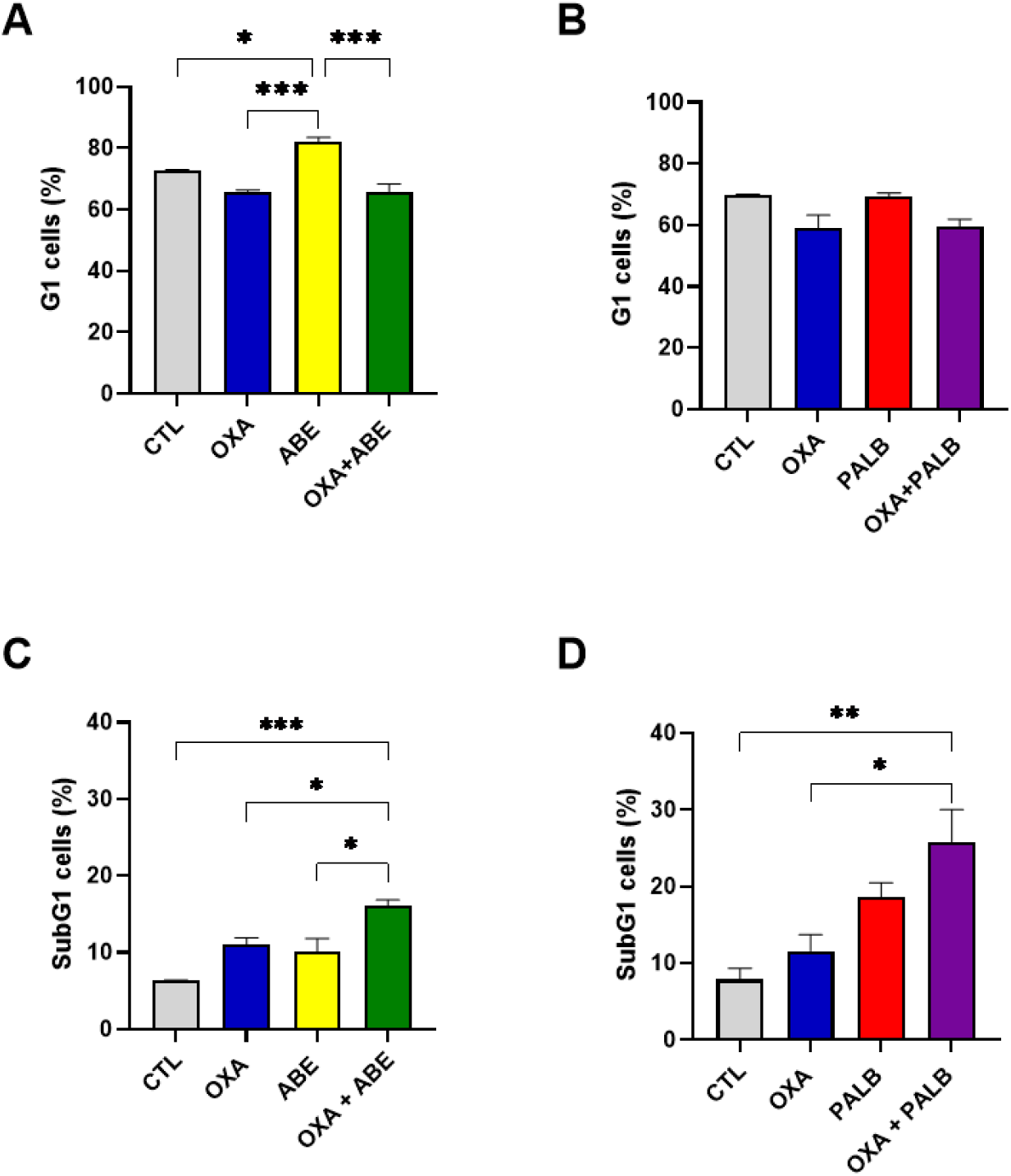
Effects of oxaliplatin combined with CDK4/6 inhibitors on cell cycle distribution and DNA fragmentation in SW480 cells. SW480 cells were treated for 48 h with oxaliplatin (1.5 μM), abemaciclib (191 nM), palbociclib (95 nM), or their respective combinations. (A–B) Percentage of cells in G₁ phase determined by flow cytometry following treatment with oxaliplatin combined with abemaciclib (A) or palbociclib (B). (C–D) Percentage of SubG₁ population, indicative of DNA fragmentation, following treatment with oxaliplatin combined with abemaciclib (C) or palbociclib (D). The data are presented as the mean ± SEM of three independent biological replicates. Statistical analysis was performed using one-way ANOVA followed by Šídák’s multiple comparisons test. Significance levels are indicated as follows: *p < 0.05; **p < 0.01; ***p < 0.001; ****p < 0.0001.

### The combination does not significantly potentiate cytotoxicity in non-tumoral IEC-6 cells

To evaluate whether the combinatorial effect is selective for tumor cells, we assessed the response of the non-tumoral intestinal epithelial cell line IEC-6 to the same treatment conditions used for HCT116 cells. IEC-6 cells were treated for 48 h with oxaliplatin (0.6 µM), abemaciclib (300 nM), palbociclib (400 nM), or their respective combinations (Fig. 9). Representative fluorescence images show viable cells (calcein, green), dead cells (ethidium homodimer, orange), and nuclei (Hoechst, blue) (Fig. 9A).

**Figure 9.**
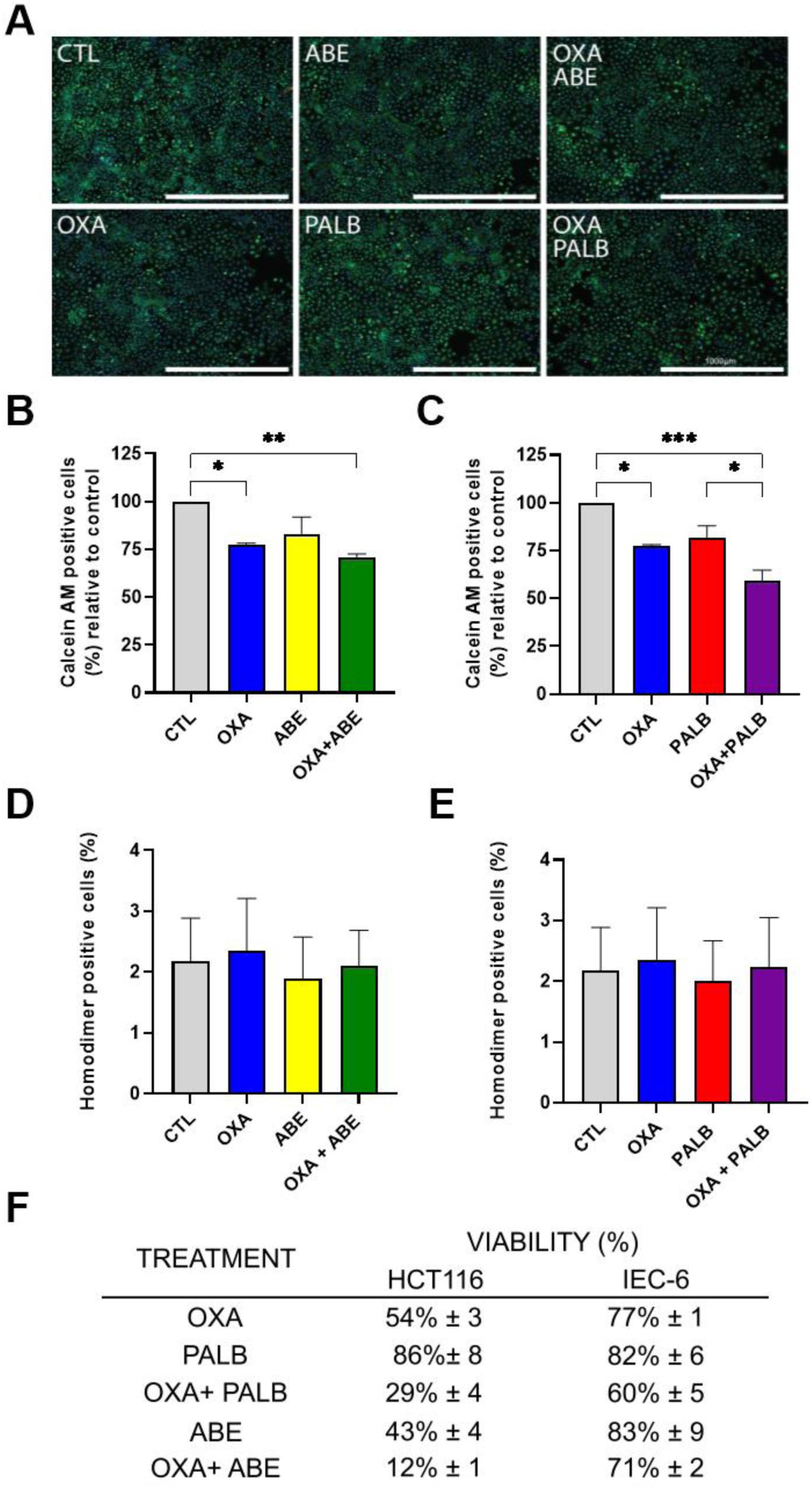
Differential effects of oxaliplatin and CDK4/6 inhibitors on cell viability in tumor and non-tumor cell lines. (A) Representative fluorescence images of IEC-6 cells treated for 48 h with oxaliplatin (0.6 μM), abemaciclib (300 nM), palbociclib (400 nM), or their combinations. Viable cells were detected by Calcein AM fluorescence (green), and nuclei were counterstained with Hoechst (blue). Scale bar: 1000 μm. (B) Quantification of Calcein AM–positive cells following treatment with oxaliplatin and abemaciclib. (C) Quantification of Calcein AM–positive cells following treatment with oxaliplatin and palbociclib. (D) Quantification of ethidium homodimer–positive cells in IEC-6 cultures treated with oxaliplatin and abemaciclib. (E) Quantification of ethidium homodimer–positive cells in IEC-6 cultures treated with oxaliplatin and palbociclib. (F) A summary table showing the percentage of viable cells in HCT116 (tumor) and IEC-6 (non-tumor) cell lines across treatments. The data are presented as the mean ± SEM of three independent biological replicates. Quantification was performed by counting Calcein AM–positive cells relative to total nuclei. Statistical analysis was performed using one-way ANOVA followed by Šídák’s multiple comparisons test. Significance levels are indicated as follows: *p < 0.05; **p < 0.01; ***p < 0.001.

Quantification of cell viability revealed that while oxaliplatin alone and the combination treatments reduced viability in IEC-6 cells relative to the untreated control, the combination of oxaliplatin with either CDK4/6 inhibitor did not significantly potentiate the effect of oxaliplatin monotherapy (Fig. 9B, C). This contrasts with the pronounced enhancement observed in HCT116 cells under identical conditions. Furthermore, assessment of cell death based on the percentage of ethidium homodimer-positive cells showed that neither monotherapies nor combination treatments increased the proportion of dead cells (Fig. 9D, E). A comparative summary (Fig. 9F) illustrates this differential response: in HCT116 cells, the combination of oxaliplatin with abemaciclib reduced viability to approximately 12%, while in IEC-6 cells, viability remained at approximately 71% under the same treatment. These findings indicate that the synergistic interaction between oxaliplatin and CDK4/6 inhibitors is preferentially active against tumor cells, supporting a tumor-selective therapeutic profile.

## Discussion

Three findings from this study are of particular translational relevance. First, among the first-line CRC chemotherapeutics tested, oxaliplatin is the most consistently synergistic partner for CDK4/6 inhibition — a conclusion supported by quantitative CI analysis across multiple concentrations and two CDK4/6 inhibitors. Second, the combinatorial effect is RB-dependent, as demonstrated by direct genetic evidence via shRNA-mediated RB1 silencing, complementing the mechanistic and correlative evidence previously reported by others [21–23]. Third, the synergy is preferentially active against tumor cells, with no significant potentiation of oxaliplatin toxicity observed in non-tumoral IEC-6 intestinal epithelial cells. Together, these findings establish a preclinical rationale for CDK4/6 inhibitor–oxaliplatin combinations in RB-intact CRC and support the existence of a tumor-selective therapeutic window.

Among the chemotherapeutic agents tested, oxaliplatin consistently exhibited the highest degree of synergism with both CDK4/6 inhibitors, whereas combinations with SN-38 yielded variable results, and 5-FU combinations approached additivity in some conditions. This differential synergistic profile may be related to the distinct mechanisms of action of these agents. CDK4/6 inhibitors arrest cells in G1, preventing entry into the S phase. SN-38, the active metabolite of irinotecan, acts primarily as a topoisomerase I inhibitor during S phase, and 5-FU is an antimetabolite that disrupts DNA and RNA synthesis predominantly in cycling cells. Both agents, therefore, necessitate active cell cycle progression for optimal cytotoxicity, which CDK4/6-induced G1 arrest may partially compromise. In contrast, oxaliplatin forms DNA interstrand crosslinks that can occur independently of cell cycle phase, and the resulting DNA damage may be amplified when cells are simultaneously prevented from engaging repair mechanisms that require S-phase entry [23]. This mechanistic compatibility may underlie the strong synergy observed between oxaliplatin and CDK4/6 inhibitors. Consistent with this interpretation, Yu et al. [23] demonstrated that CDK4/6 inhibition enforces RB1/TEAD4/HDAC1-mediated epigenetic suppression of DNA repair genes, thereby sensitizing CRC cells to oxaliplatin-induced DNA damage. The rationale for combining cell cycle-targeted agents with conventional chemotherapy to exploit complementary resistance vulnerabilities has been previously discussed [30].

Our data reveal functional differences between abemaciclib and palbociclib in the context of combination therapy with oxaliplatin. In HCT116 cells, abemaciclib induced a more robust G1 arrest as a single agent, and its combination with oxaliplatin significantly increased the sub-G1 population, indicative of enhanced cell death. In contrast, palbociclib combined with oxaliplatin did not significantly increase the sub-G1 fraction in this cell line, despite reducing overall viability. Importantly, both inhibitors reduced pRB phosphorylation at Ser807/811 by approximately 50% in HCT116 cells, suggesting that the functional differences between the two agents are unlikely to be explained solely by differential target engagement at the level of pRB. Abemaciclib is known to possess broader kinase inhibitory activity compared to palbociclib, including activity against CDK9, GSK3α/β, and other kinases [31, 32], which may contribute to its enhanced capacity to induce cell death beyond pure cytostasis in this cellular context. Additionally, abemaciclib has been shown to accumulate in lysosomal compartments, a phenomenon known as "lysosomal trapping," which may promote functional and morphological alterations associated with atypical cell death mechanisms [33, 34]. However, this apparent superiority of abemaciclib in promoting sub-G1 accumulation was not observed in SW480 cells, where the combination of oxaliplatin with palbociclib produced a larger sub-G1 increase than the abemaciclib combination. In summary, these results indicate that although abemaciclib’s wider kinase inhibitory profile may be beneficial in some CRC situations, it is not possible to generalize inhibitor selection based on anticipated sub-G1 behavior across tumor models without taking cellular context into account. Interestingly, the most pronounced cell death responses were observed in conditions where G1 arrest was not further enhanced by the combination, suggesting that the relative contribution of cytostatic versus cytotoxic outcomes may depend on the cellular context and the inhibitor used—a question that warrants further investigation.

RB1 silencing in HCT116 cells partially attenuated the viability reduction induced by the combination of oxaliplatin with either CDK4/6 inhibitor, supporting the hypothesis that the combinatorial effect is RB-modulated. This finding is consistent with the established role of RB as the primary mediator of CDK4/6 inhibitor activity [17, 35] and aligns with recent evidence that FOLFOX-resistant CRC cells exhibit marked reduction of RB and phospho-RB protein levels, suggesting RB status as a determinant of therapeutic response [21]. Of note, Schneider et al. [21] evaluated all three clinically approved CDK4/6 inhibitors—ribociclib, palbociclib, and abemaciclib—in a broad panel of CRC cell lines and demonstrated that high p16 expression correlates with resistance across all three agents; however, their combination experiments with FOLFOX were performed exclusively with ribociclib and did not reveal synergism in that specific context. Our research builds on these findings by offering direct genetic evidence, via shRNA-mediated RB1 knockdown, that the synergistic interaction between oxaliplatin and CDK4/6 inhibitors is influenced by RB expression.

Beyond its canonical role in controlling G1/S transition, the CDK4/6–RB axis has been implicated in cellular differentiation, genomic stability, regulation of programmed cell death, autophagy, and senescence [36–40]. Although the latter is a recognized outcome of CDK4/6 in the context of inhibitor treatment, the 48-hour treatment window used in this study does not allow for the establishment of a stable senescent phenotype, which typically requires several days of continuous exposure, as shown by studies using 5-day protocols to detect senescence-associated β-galactosidase activity in CDK4/6 inhibitor-treated tumor cells [41]. The predominance of G1 arrest, sub-G1 accumulation, and increased ethidium homodimer staining observed at this timepoint collectively indicate that cytostatic and cytotoxic responses are the primary cellular outcomes under our experimental conditions. In addition, aberrant pRB hyperphosphorylation has also been reported in the colonic epithelium of patients with Crohn’s disease [42], an inflammatory condition associated with increased CRC risk, further underscoring that dysregulation of this axis in the intestinal epithelium extends beyond established neoplasia. The engagement of these non-canonical functions may contribute to the complexity of the combinatorial response observed in our study.

A notable finding of our study is that synergistic benefit was maintained even when abemaciclib was used at sub-cytotoxic concentrations (IC₂₀) in combination with oxaliplatin in SW480 cells for both CDK4/6 inhibitors. This observation could have a significant translational implication. CDK4/6 inhibitors are associated with dose-dependent adverse effects, most notably neutropenia, and represent a substantially higher cost relative to conventional chemotherapeutics such as oxaliplatin. The finding that dose reduction of the targeted agent preserves synergistic efficacy suggests that lower-dose combination regimens could be clinically feasible, potentially improving tolerability and expanding access in resource-limited settings, aligning with the broader concept that strategic combinations of novel targeted therapies with established chemotherapeutics can enhance tumor cell death while minimizing toxicity [43]. This idea is backed up by the clinical use of trilaciclib, a CDK4/6 inhibitor given in low doses before chemotherapy to protect hematopoietic cells and boost antitumor immunity at the same time [24]. This shows that CDK4/6 inhibition can have a real clinical benefit even at doses lower than the maximum tolerated dose.

The combinatorial effect of CDK4/6 inhibitors with oxaliplatin was confirmed in SW480 cells, where both abemaciclib and palbociclib enhanced oxaliplatin-induced sub-G1 accumulation at sub-cytotoxic inhibitor concentrations. However, the effect required higher concentrations of oxaliplatin and CDK4/6 inhibitors. This difference may be influenced by several genetic and epigenetic distinctions between the two cell lines, including, among other factors, the distinct TP53 status: HCT116 harbors wild-type TP53, whereas SW480 carries the inactivating R273H mutation [44, 45]. Loss of functional p53 compromises DNA damage-induced apoptotic signaling, which may limit the additional cytotoxic benefit conferred by CDK4/6 inhibition in the context of oxaliplatin-induced DNA damage [46]. Furthermore, p53 signaling within the tumor microenvironment has been shown to contribute to tissue chemoresistance through multiple mechanisms beyond cell-autonomous apoptosis [47]. Nonetheless, the observation that CDK4/6 inhibitors potentiated oxaliplatin-induced cell death even in this less sensitive cellular background, and at sub-cytotoxic inhibitor doses, reinforces the broader applicability of this combinatorial strategy across CRC contexts.

The combination of oxaliplatin with CDK4/6 inhibitors did not significantly potentiate the effect of oxaliplatin monotherapy in non-tumoral IEC-6 intestinal epithelial cells, in contrast to the pronounced synergistic enhancement observed in HCT116 tumor cells. This differential response suggests a potential therapeutic window favoring tumor cells, which is particularly relevant given the gastrointestinal toxicity associated with platinum-based regimens [6]. The preservation of IEC-6 viability is consistent with evidence that CDK4/6 inhibition can protect normal tissues from chemotherapy-induced damage through RB-dependent mechanisms [48–50]. In RB-intact non-tumoral cells, CDK4/6-induced G1 arrest may serve a protective function by preventing cells from entering S phase while the genotoxic insult is present, thereby reducing vulnerability to DNA damage. Conversely, in tumor cells with hyperactivated CDK4/6 signaling, pharmacological inhibition disrupts an oncogenic dependency, amplifying rather than attenuating the cytotoxic effect of oxaliplatin.

Several aspects of the present study address gaps in the current literature. To our knowledge, this is the first systematic head-to-head comparison of three first-line CRC chemotherapeutic agents as combinatorial partners for CDK4/6 inhibition, using quantitative synergism analysis (Chou-Talalay). While previous studies have examined individual drug pairs—palbociclib with irinotecan [22] or CDK4/6 inhibition with oxaliplatin [23]—our approach identifies oxaliplatin as the optimal partner through direct comparison within the same experimental system. Additionally, the parallel evaluation of two CDK4/6 inhibitors reveals functional distinctions between abemaciclib and palbociclib that are relevant for clinical drug selection. The use of shRNA-mediated RB1 knockdown provides direct genetic evidence of RB dependency in the combinatorial context, complementing the pharmacological and correlative evidence reported by others [21, 23]. Finally, the assessment of tumor selectivity using a non-tumoral intestinal epithelial model addresses a clinically important question that has not been previously examined for this drug combination in CRC.

Our findings were obtained in two-dimensional monolayer models, which enabled systematic pharmacological characterization but do not fully recapitulate the complexity of the tumor microenvironment. The assessment of tumor selectivity using IEC-6 cells, a well-established non-tumoral intestinal epithelial model previously employed to characterize differential responses to CDK inhibitor treatment in the intestinal context [51], provides an initial indication of a favorable therapeutic index for this drug combination. Future studies employing three-dimensional culture systems, including microfluidic tumor-on-a-chip platforms that generate physiologically relevant nutrient and drug gradients [52], will be important to further substantiate these findings, define the scope of tumor selectivity, and evaluate the translational potential of this combination strategy.

In conclusion, our findings demonstrate that oxaliplatin combined with CDK4/6 inhibitors produces synergistic antitumor effects in CRC cells through an RB-dependent mechanism involving sustained G1 arrest and enhanced cell death. The combinatorial benefit is maintained at sub-cytotoxic inhibitor concentrations and is preferentially active against tumor cells, supporting a favorable therapeutic index. These results provide a preclinical rationale for the continued investigation of oxaliplatin–CDK4/6 inhibitor combinations in CRC, with attention to RB status as a potential biomarker for patient stratification.

## Conflict of interest statement

The authors declare no conflict of interest.

## Acknowledgments

This study was supported by the Programa Nacional de Apoio à Atenção Oncológica (PRONON/Ministério da Saúde, Lei n° 12.715/2012, NUP 25000.016131/2018-77); the Fundação Carlos Chagas Filho de Amparo à Pesquisa do Estado do Rio de Janeiro (FAPERJ), grant nos. E-26/210.201/2022, E-26/210.623/2023, and 260003/001162/2023; the Conselho Nacional de Desenvolvimento Científico e Tecnológico (CNPq), grant no. 308046/2025-0; and the Coordenação de Aperfeiçoamento de Pessoal de Nível Superior — Brasil (CAPES) — Finance Code 001. The authors thank Prof. José Garcia Abreu (ICB/UFRJ) for kindly providing the IEC-6 cell line and Daniel Cadilhe for administrative support.

## Author Contributions

**A.S.O.S.:** Investigation, Formal analysis, Visualization, Writing – original draft. **J.S.M.C.:** Investigation, Formal analysis, Visualization, Writing – original draft. **L.S.F.:** Investigation, Formal analysis, Visualization, Writing – original draft. **J.P.A.D.:** Formal analysis, Visualization, Writing – review & editing. **B.R.M.:** Investigation, Writing – review & editing. **C.V.:** Investigation, Writing – review & editing. **M.S.C.:** Methodology, Writing – review & editing. **S.K.R.:** Resources, Funding acquisition, Writing – review & editing. **M.H.B.:** Methodology, Resources, Writing – review & editing. **H.L.B.:** Conceptualization, Supervision, Project administration, Funding acquisition, Writing – review & editing.

## Declaration of generative AI and AI-assisted technologies in the manuscript preparation process

During the preparation of this work the author(s) Claude Opus 4.6 (Anthropic, 2026) was used to assist with language editing and manuscript preparation. All content was critically reviewed and approved by the authors, who take full responsibility for the work.

## Notes

### Competing Interest Statement

The authors have declared no competing interest.

